# Diverse food-sensing neurons trigger idiothetic local search in *Drosophila*

**DOI:** 10.1101/433771

**Authors:** Román A. Corfas, Michael H. Dickinson

## Abstract

Resources are often sparsely clustered in nature. Thus, foraging animals may benefit from remembering the location of a newly discovered food patch while continuing to explore nearby [1, 2]. For example, after encountering a drop of yeast or sugar, hungry flies often perform a local search consisting of frequent departures and returns to the food site [3, 4]. Fruit flies, *Drosophila melanogaster*, can perform this food-centered search behavior in the absence of external stimuli or landmarks, instead relying solely on internal (idiothetic) cues to keep track of their location [5]. This path integration behavior may represent a deeply conserved navigational capacity in insects [6, 7], but the neural pathways underlying food-triggered searches remain unknown. Here, we used optogenetic activation to screen candidate cell classes and found that local searches can be initiated by diverse sensory neurons including sugar-sensors, water-sensors, olfactory-receptor neurons, as well as hunger-signaling neurons of the central nervous system. Optogenetically-induced searches resemble those triggered by actual food and are modulated by starvation state. Furthermore, search trajectories exhibit key features of path integration: searches remain tightly centered around the fictive-food site, even during long periods without reinforcement, and flies re-center their searches when they encounter a new fictive-food site. Flies can even perform elaborate local searches within a constrained maze. Together, these results suggest that flies enact local searches in response to a wide variety of food-associated cues, and that these sensory pathways may converge upon a common neural system for path integration. Optogenetically induced local searches in *Drosophila* can now serve as a tractable system for the study of spatial memory and navigation in insects.

## RESULTS & DISCUSSION

### Multiple sensory pathways trigger local search

To discover sensory pathways triggering local search, we tracked the behavior of individual, food-deprived female flies as they explored a circular arena with a featureless optogenetic activation zone at its center (Figure 1A). The assay consists of an initial 10-minute baseline control period, followed by a 30-minute period during which animals receive a 1-second pulse of red light (628 nm) whenever they enter the activation zone. For flies expressing the light-sensitive channel *CsChrimson* in food-sensing neurons, the activation zone should act as a patch of fictive food, potentially able to elicit a local search. Aside from the light pulses used for optogenetic activation, the animals are in complete darkness and must rely on internal cues to navigate the open-field portion of the arena.

**Figure 1.**
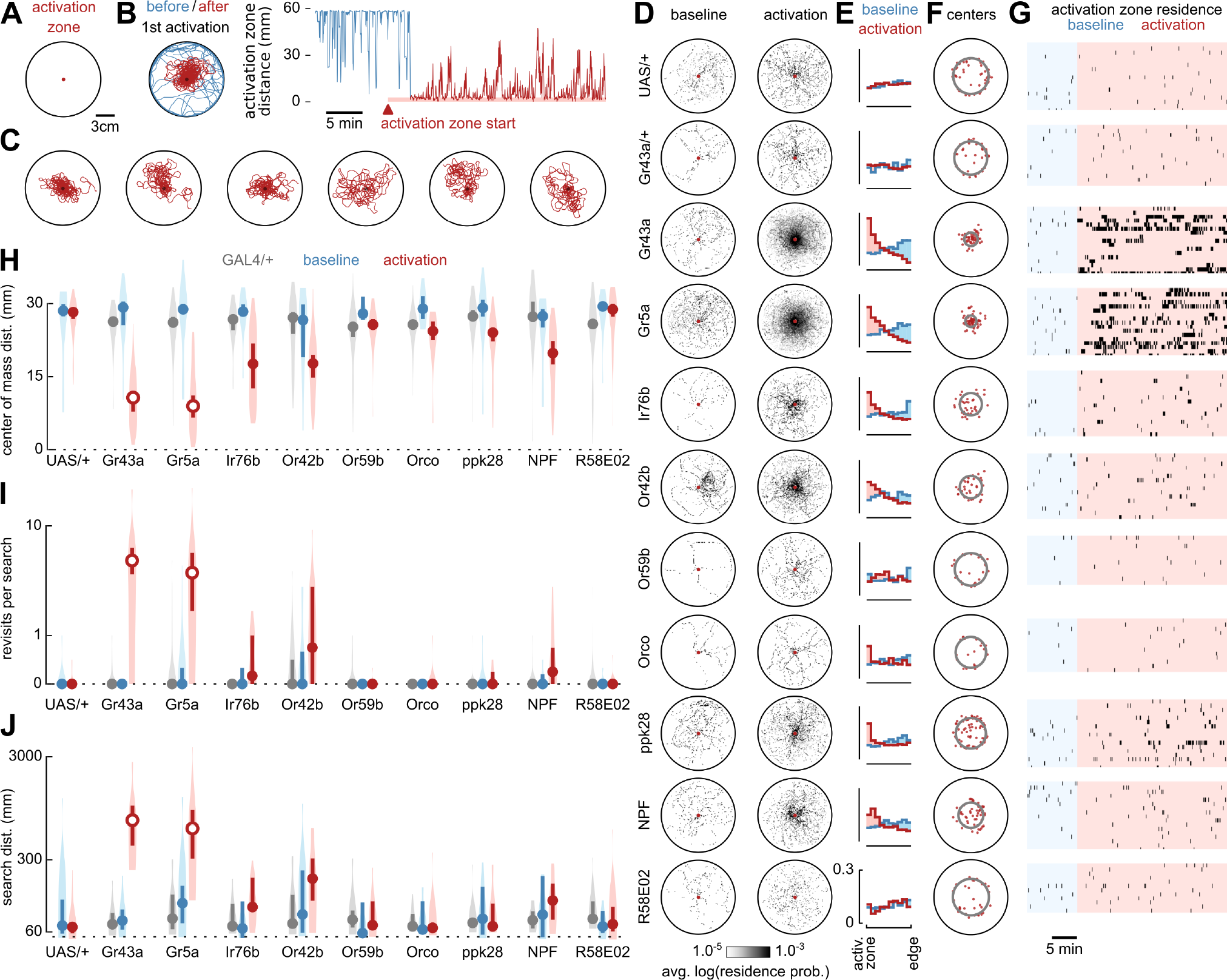
Optogenetic activation of food-sensing neurons triggers local search. (A) Schematic of experimental arena featuring an optogenetic activation zone. A female *Drosophila* is about 3 mm in length. (B) Left: example trajectory of *Gr43a-GAL4>UAS-CsChrimson* fly before (blue) and after (red) first optogenetic stimulation. Right: the same data, plotted as the distance between the fly and activation zone center. The activation zone becomes operational after an initial 10-minute baseline control period. (C) Six of the seven longest distance search bouts (left to right) triggered by *Gr43a-GAL4* activation; the longest bout is plotted in (B). Data include search bouts truncated by the conclusion of the trial. (D) Residence probabilities of walking trajectories during activation search bouts (with light pulses) and baseline control bouts (without light pulses). Each spatial probability histogram shows the mean of the mean normalized residence probability distribution of each fly. Throughout the paper (unless otherwise noted), analyses excluded search bouts truncated by the conclusion of the trial, as well as any timepoints when the fly was stopped or in the activation zone. UAS/+ indicates data for *UAS-CsChrimson*;+ parental controls. Gr43a/+ (for example) indicates data for *Gr43a-GAL4*;+ parental controls whereas Gr43a indicates corresponding data for *Gr43a-GAL4>UAS-CsChrimson* experimental flies. (E) Probability distributions of distance between fly and activation zone during activation search bouts (red) or baseline control bouts (blue). Each probability histogram shows the mean of the mean normalized probability distribution of each fly. (F) Centers of mass for all activation search bouts (red dots). Grey rings show the distance from the activation zone of the median center of mass. (G) Raster plots of activation zone residence during baseline (blue) and while the activation zone is operational (red). Marks are graphically extended horizontally by 5 seconds for visibility. (H) Distance from the arena center for centers of mass of baseline control bouts (blue), activation search bouts (red) and activation search bouts of GAL4-line parental controls (grey). Circles depict medians, error bars depict 95% confidence intervals, and violin plots indicate full data distribution. Unfilled circles indicate cases in which the median is statistically different (p≤0.05, with Bonferroni correction) from both baseline condition (Wilcoxon signed-rank test) and GAL4-line parental control (where applicable, Mann-Whitney U test). (I) Mean number of revisits to the activation zone (plotted on a log axis) during baseline control bouts (blue), activation search bouts (red), and activation search bouts of GAL4-line parental controls (grey). Plotting conventions as in (H). (J) Mean distance walked (plotted on a log axis) during baseline control bouts (blue), activation search bouts (red), and activation search bouts of GAL4-line parental controls (grey). Plotting conventions as in (H).

It is known that flies perform local searches after discovering a drop of sucrose [3–5, 8–10], suggesting that sweet-sensing neurons may be sufficient to initiate this behavior. To test this, we used *Gr43a-GAL4>UAS-CsChrimson* flies to activate fructose-sensing neurons whenever the flies entered the activation zone [11, 12]. Activation of these gustatory neurons triggered local searches remarkably similar to those previously observed in response to actual food (n=20 flies; Figures 1B and 1C; Movies 1 and 2) [5], indicating that this sensory pathway is sufficient to elicit a sustained bout of idiothetic path integration. Unlike parental controls (n=15-23 flies), *Gr43a-GAL4>UAS-CsChrimson* flies extensively searched the area surrounding the activation zone (~30 cm^2^), after receiving a light-pulse (Figures 1D and 1E). These search trajectories were highly centered at the activation zone (Figures 1F and 1H) and consisted of numerous revisits to the activation zone (Figures 1G, 1I, and S1A-S1C) — both features of local searches shown to require path integration [5]. During local search, flies cumulatively walked ~30-300 cm before eventually straying to the arena edge (Figure 1J). Nearly identical local searches were triggered by activation of sugar-sensing neurons using the *Gr5a-GAL4* driver (n=20 flies; Figures 1D-1J, S2A-S2D; Movie 3) [13, 14]. This result demonstrates that non-pharyngeal sugar sensors are capable of eliciting local search, in contrast to recent experiments suggesting otherwise [10]. The use of fictive food in these experiments provides further evidence that flies are in fact using idiothetic path integration during local search, rather than relying on external (allothetic) cues coming from an actual drop of food such as visual features, odor or humidity gradients, or tracks of food residue deposited during search excursions.

Prior research has shown that, compared to sucrose-triggered searches, a drop of 5% yeast solution elicits search trajectories that are even longer and include more revisits to the food [5, 15], suggesting that proteinaceous food components may also have a role in initiating this behavior. This would make sense, given that amino acids present in yeast are a coveted source of nutrition for mated females, which require a protein source to produce eggs [16–18]. The ionotropic receptor Ir76b has been implicated in the detection of amino acids [19, 20] and other important nutrients such as salt [21], polyamines [22], and fatty acids [23]. We tested *Ir76b-GAL4>UAS-CsChrimson* flies in our assay, and found that activation of these amino-acid sensors produced a much less extensive local search than that elicited by sweet-sensing neurons. The trajectories were less centered at the activation zone, covered less distance, and rarely included a revisit to the activation site (n=16 flies; Figures 1D-1J and S2A-S2D).

Food odorants also trigger search behavior in insects. In flight, for example, encounters with an odor plume elicit the stereotyped cast and surge maneuvers that enable insects to localize the source of an advected odor [24, 25]. Recent studies have demonstrated that this also occurs during walking—flies increase their turn rate when they exit a plume of apple cider vinegar (ACV) odor [26, 27]. Attraction to the smell of ACV in *Drosophila* is mediated primarily by neurons expressing the olfactory receptor *Or42b* [28]. Optogenetic activation of *Or42b-GAL4* neurons produces attraction behavior in flies [29], as does activation of *Or59b-GAL4* neurons [30], which respond to acetate esters found in food odors [31, 32]. Simultaneous optogenetic activation of nearly all the olfactory receptor neurons via *Orco-GAL4* also produces attraction in flies [29]. We tested whether these three classes of olfactory neurons could trigger a local search and found that activation of *Orco*- (n=13 flies) and *Or59b-GAL4* (n=12 flies) neurons did not elicit searches (Figures 1D-1J and S2A-S2D). In contrast, activation of ACV-odor-sensing *Or42b-GAL4* neurons triggered local searches; although these were modest in comparison to those triggered by sugar-sensing neurons, they nevertheless covered hundreds of millimeters and typically featured at least one revisit to the activation zone (n=16 flies; Figures 1D-1J and S2A-S2D).

Next, we tested whether the water content of food drops might be enough to evoke local search. In *Drosophila*, water sensation is mediated by the osmosensitive ion channel ppk28, a member of the degenerin/epithelial gene family [33]. We found that activation of water-sensing *ppk28-GAL4* neurons in food-deprived flies resulted in a modest increase in residence near the activation zone (n=18 flies; Figures 1D and 1E), due largely to the animals ceasing locomotor activity (see results below); however, the activation did not trigger a local search (Figures 1F-1J and S2A-S2D). This result is in agreement with previous behavioral experiments showing that *Drosophila* do not produce local search bouts after encountering a drop of pure water [5, 9].

Because we found that a variety of fictive food stimuli evoke local searches, we hypothesized that reward-signaling neurons of the central nervous system might also trigger the behavior. In other words, flies might initiate local searches around any location associated with a rewarding stimulus, even without accompanying activation of peripheral food-sensing chemosensors. To examine this possibility, we tested activation of either neuropeptide-F (NPF) or protocerebral anterior medial (PAM) neurons. NPF is a highly conserved hunger-signaling neuropeptide that stimulates a variety of *Drosophila* behaviors, including feeding [34]. *NPF-GAL4* labels neurons in the posterior region of the *Drosophila* brain, and activation of these cells is rewarding in the context of olfactory conditioning [35]. We found that activation of *NPF-GAL4* neurons results in modest local searches, similar in extent to those triggered by *Ir76b-GAL4* neurons (n=18 flies; Figures 1D-1J and S2A-S2D). Another set of reward-signaling neurons are dopaminergic PAM neurons, which are activated by sugar ingestion and innervate the mushroom body, a structure critical for forming associative memories [36, 37]. Activation of PAM neurons via *R58E02-GAL4* is known to mediate reward during olfactory conditioning [36, 37], and silencing PAM neurons inhibits food occupancy during foraging [38]. However, we found that activation of PAM neurons does not produce search behavior (n=12 flies; Figures 1D-1J and S2A-S2D).

### The extent of local search is modulated by starvation state

Because local searches are initiated by food-associated chemosensors and hunger-signaling neurons, we hypothesized that starvation state may influence the extent of optogenetically induced searches. The influence of starvation has been observed for sucrose-induced searches in *Drosophila* [9] as well as protein- and water-induced searches in the blowfly (*Phormia regina*) [15]. Until this point, all of our experiments were conducted with animals allowed access only to water for 33-42 hours preceding the trial. To examine the importance of starvation in promoting local search, we tested activation of sugar-sensing *Gr43a-GAL4* neurons in flies that were reared continually on food (n=16 flies) or starved for only 9-18 hours (n=18 flies). As expected, we found that longer starvation times result in more extensive searches, with longer trajectories and more revisits to the activation zone (Figures 2A-2D).

**Figure 2.**
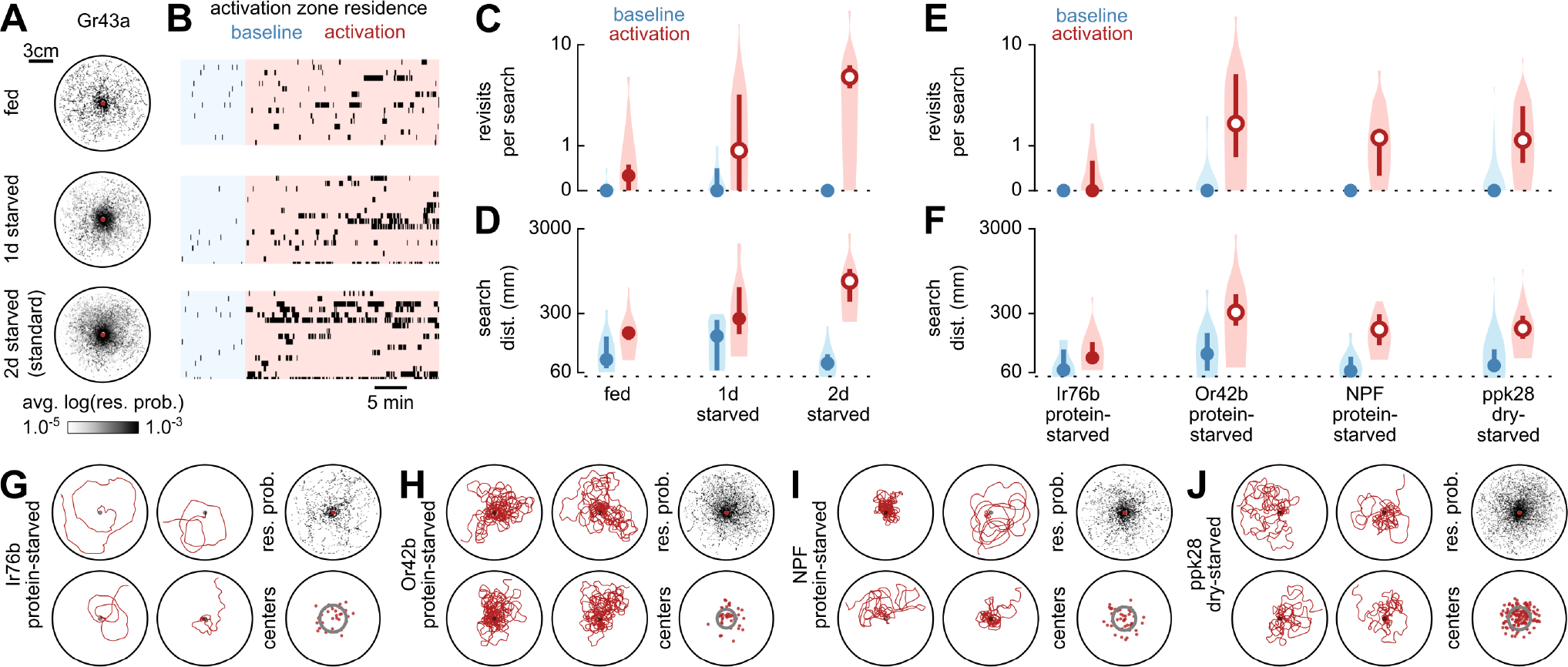
Local search around fictive food is augmented by starvation. (A) Residence probabilities during activation search bouts for *Gr43a-GAL4>UAS-CsChrimson* animals reared continually on food (fed), starved for 9-18 hours (1d starved), or starved for our standard 33-42 hours (2d starved), plotted as in Figure 1D. 2d starved data are replotted from Figures 1D-1J for convenience. (B) Raster plots of activation zone residence for data in (A), plotted as in Figure 1G. (C) Mean number of revisits to the activation zone for data in (A), plotted as in Figure 1I. (D) Mean distance walked during search bouts for data in (A), plotted as in Figure 1J. (E) Mean number of revisits to the activation zone for animals of the indicated genotype and starvation state, plotted as in Figure 1I. (F) Mean distance walked during search bouts for animals of the indicated genotype and starvation state, plotted as in Figure 1J. (G) Longest distance search bouts (left, clockwise from top-left, plotted as in Figure 1C), residence probabilities (top-right, plotted as in Figure 1D), and centers of mass (bottom-right, plotted as in Figure 1F), for activation search bouts of protein-starved *Ir76b-GAL4>UAS-CsChrimson* animals. (H) As in (G) for protein-starved *Or42b-GAL4>UAS-CsChrimson* animals. (I) As in (G) for protein-starved *NPF-GAL4>UAS-CsChrimson* animals. (J) As in (G) for dry-starved *ppk28-GAL4>UAS-CsChrimson* animals.

Protein-deprived *Drosophila* show increased local exploration of yeast patches during foraging [39]. Thus, we next tested whether protein deprivation could strengthen searches triggered by sensory pathways that elicited weak searches in our original screen. Even in animals subjected to this condition, activation of amino acid-sensing *Ir76b-GAL4* neurons did not elicit extensive local search (n=15 flies; Figures 2E-2G and S3A-S3D), despite the fact that protein-deprived mated females are known to develop a strong preference for amino acid-containing food [16–18]. However, we found that activation of ACV-odor-sensing *Or42b-GAL4* neurons in protein-starved animals resulted in more extensive and centralized local searches, now comparable to those triggered by sugar-sensing neurons (n=13 flies; Figures 2E-2F, 2H, and S3A-S3D; Movie 4). This result is consistent with work showing that starvation promotes food search behavior in *Drosophila*, and that this effect is mediated by neuropeptidergic modulation of *Or42b-GAL4* neuron activity [40]. We found a similar enhancement of local search in protein-starved *NPF-GAL4>UAS-CsChrimson* animals (n=11 flies; Figures 2E-2F, 2I, and S3A-S3D; Movie 5). Remarkably, nutritional deprivation can even produce searches triggered by water-sensation—activation of *ppk28-GAL4* neurons elicited robust local searches in animals subjected to a desiccating environment without food or water, (n=25 flies; Figures 2E-2F, 2J, and S3A-S3D; Movie 6). Collectively, these results suggest that optogenetically-induced searches are influenced by internal nutrient states in a similar way to searches triggered by actual food.

In his initial description of food-induced local search, Vincent Dethier demonstrated that when a hungry blowfly discovered the drop of food, it performed a proboscis extension response (PER)—a reflex associated with appetitive cues [3, 4]. To explore the role of proboscis extension in optogenetically induced local search, we tested whether activation of each neuron class elicits PER. As has been previously reported, activation of sugar-sensing *Gr5a-GAL4* neurons elicits PER (Figure S4A) [41–43]. We found that activation of *Gr43a-GAL4* neurons also elicits PER in a starvation-dependent manner (Figure S4A), indicating that fructose triggers a feeding reflex similar to that of other sugars. Activation of water sensors via *ppk28-GAL4* neurons also resulted in PER, even in animals that had not been subjected to dry-starvation (Figure S4A). We also found that activation of hunger-signaling neurons via *NPF-GAL4* elicited strong PER (Figure S4A), demonstrating a novel function for these neurons. However, none of the other neuron classes in our screen consistently triggered PER, including *Or42b-GAL4* neurons (Figure S4A), indicating that local search can be initiated by receptors that do not by themselves elicit PER.

Together, these results suggest that local searches are triggered by both contact chemosensory cues that signal that the fly is on food (e.g. water or sugar), as well as volatile cues that indicate that the fly is in the vicinity of food (e.g. the odor of ACV). Although searches triggered by sugar, water, and odor sensation appear broadly similar in our experiments (Figures 1C, 2H and 2J), it is likely that their underlying behavioral structure differs [26, 44]. For example, we found that whereas activation of *Gr43a-*, *Gr5a-Ir76b-*, *ppk28-*, or *NPF-GAL4* results in a drop in locomotor rate or complete stopping, activation of ACV-odor-sensing *Or42b-GAL4* neurons only elicits a brief startle response similar to controls (Figures S4B and S4C). The absence of slowing at the initiation of searches triggered by *Or42b-GAL4* neurons is consistent with the interpretation that these searches are related to the casting behaviors elicited by loss of an odor plume [24, 25, 27]. Additional studies will be necessary to determine whether local searches triggered by food-associated stimuli are stereotyped, or are instead accomplished through diverse behavioral mechanisms.

### Optogenetically-induced local searches provide a model system for the study of insect path integration

Our results show that optogenetically-induced local searches resemble those evoked by actual food, suggesting that flies are using idiothetic path integration to keep track of their position relative to the activation zone. Unlike previous studies using real food [3–5, 8–10], we are able to monitor every occasion that the fly senses the fictive food and therefore has an opportunity to reinforce or reset the memory of the stimulus location. Analyzing data from our screen, we found that *Gr43a-GAL4>UAS-CsChrimson* animals can perform a centered local search lasting minutes and covering hundreds of millimeters without an intervening optogenetic stimulation (Figures 3A-3C). This implies that a persistent internal representation of space underlies this behavior. We also observed that flies can update the center of their search upon discovering another activation zone, as has been found with searches around real food [5], and moreover that flies can repeatedly shift the center of their search between activation zones (n=13 flies; Figures 3D-3F).

**Figure 3.**
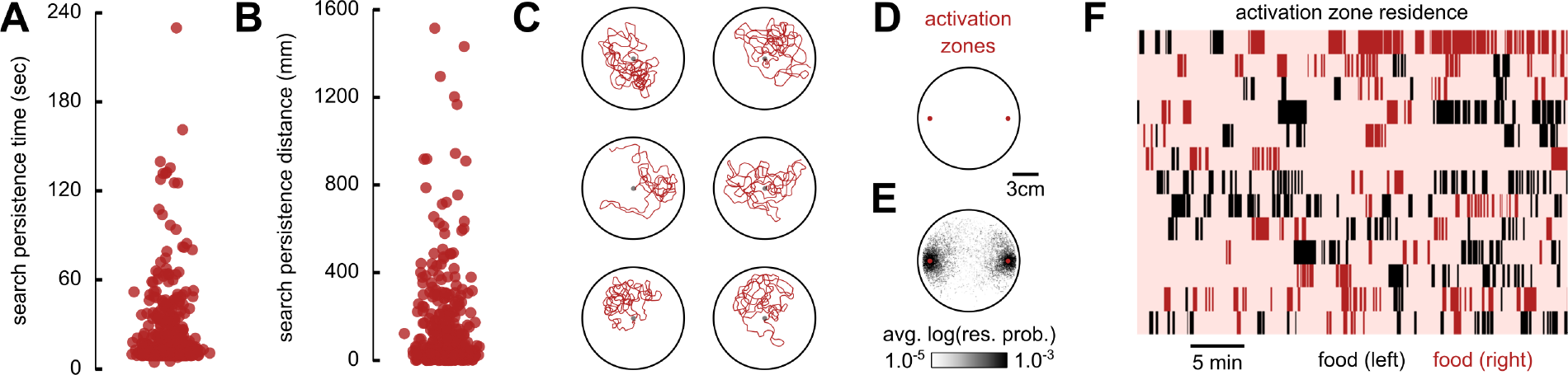
Flies keep track of fictive-food location during optogenetically-induced local search. (A) Duration of all trajectories beginning at the offset of a light pulse and ending with onset of the subsequent light pulse or conclusion of search bout, for *Gr43a-GAL4>UAS-CsChrimson* animals. Data include residence in the activation zone and search bouts truncated by the conclusion of the trial. (B) Distance walked for each trajectory plotted in (A). (C) Longest distance trajectories (clockwise from top-left) from data in (B). (D) Schematic of experimental arena with two optogenetic activation zones. (E) Residence probability during activation search bouts, plotted as in Figure 1D. (F) Raster plots of residence in both activation zones, plotted as in Figure 1G.

To explore the versatility of local search behavior, we investigated whether flies can execute path integration in a constrained environment, as opposed to an open field. We constructed a grid-shaped maze called flyadelphia—a reference to Philadelphia’s canonical street grid—with an optogenetic activation zone at its center (Figure 4A). Remarkably, *Gr43a-GAL4>UAS-CsChrimson* animals were able to perform lengthy and elaborate local searches within this confined arena (n=16 flies; Figure 4B; Movie 7). In the absence of a light-pulse, flies typically walk through the activation zone and continue in a straight path until reaching the arena edge (n=15 flies; Figures 4C-G). In contrast, when subjected to a light pulse, flies explore the square blocks surrounding the activation zone, covering large search distances and frequently revisiting the activation zone (Figures 4C-G). These experiments demonstrate that flies can execute a local search without being able to freely choose the location, timing, or angle of their turns—further evidence that this behavior relies on path integration rather than just a random search process [5].

**Figure 4.**
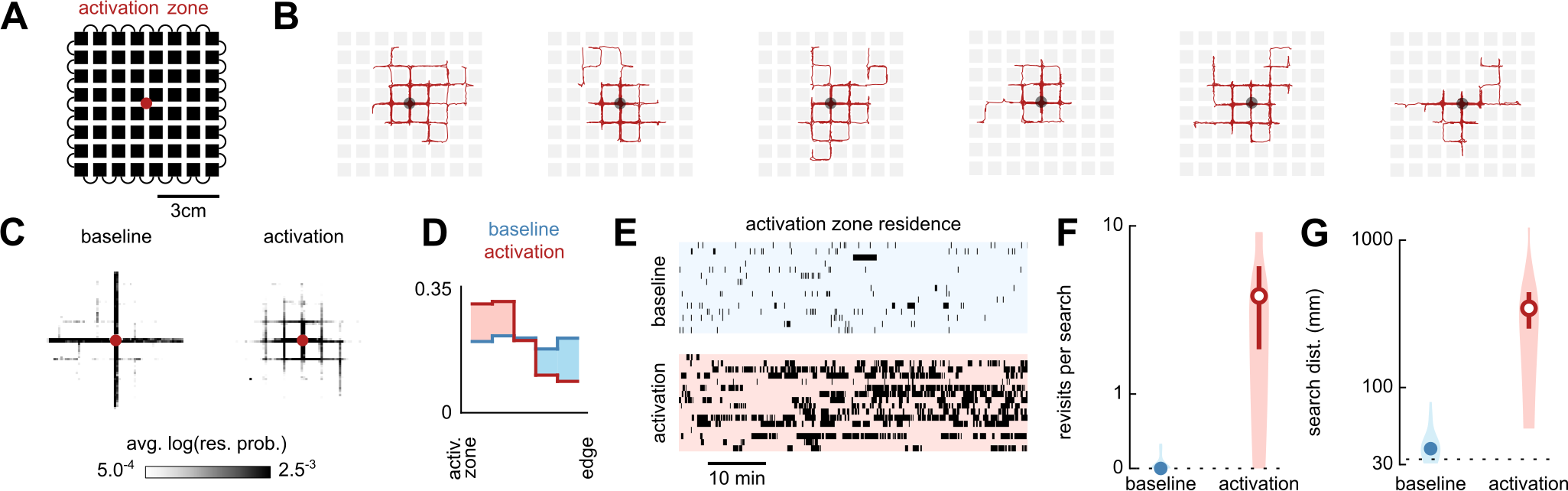
Flies perform local search in a confined maze. (A) Schematic of flyadelphia experimental arena featuring an optogenetic activation zone. Flies are restricted to the narrow passages between the blocks of the grid (black). (B) Longest distance search bouts (from left to right, plotted as in Figure 1C). (C) Residence probability during search bouts, plotted as in Figure 1D. (D) Probability distributions of fly distance to the activation zone, plotted as in Figure 1E. (E) Raster plots of activation zone residence during baseline (top) or activation (bottom) experiments, plotted as in Figure 1G. (F) Mean number of revisits to the activation zone during search bouts, plotted as in Figure 1I. (G) Mean distance walked during search bouts, plotted as in Figure 1J.

The ability to execute a sustained search centered around a fictive-food site in complete darkness, and moreover to carry this out in an environment with arbitrary geometric constraints, strongly suggests that flies can keep track of their location relative to the activation zone. This feat of idiothetic path integration has previously been compared to other insect behaviors such as the foraging excursions of desert ants (*Cataglyphis fortis*) [5], which routinely embark on long and winding runs through featureless terrain and yet are able to return to their nest in a direct path [45]. To perform dead-reckoning, these ants keep track of both the distance and the direction of their travel, enabling them to integrate their position relative to a point of origin [46–48]. During food-triggered searches, *Drosophila* may be using the same computational strategies as *Cataglyphis*, and thus may be relying on the same highly conserved brain structures [6, 7]. In particular, studies point to the importance of the central complex—a sensorimotor hub of the insect brain that processes numerous aspects of locomotion, navigation and decision-making [49]. Wedge neurons of the ellipsoid body encode azimuthal heading, potentially serving as a compass for path integration, celestial navigation, and other behaviors [50–53]. Whereas less is known about how insects monitor odometry, it is thought that step-counting can be achieved by using proprioceptive feedback or efferent copies of motor commands to integrate distance traveled [6, 47].

We propose that optogenetic activation of *Gr43a-* and *Gr5a-GAL4* sugar sensors may be a potent tool in future experiments seeking to characterize the neural implementation of path integration. Among the sensory pathways we studied, these sweet-sensing neurons are the most reliable triggers of local search. However, the comparatively weaker searches elicited by activation of other neural pathways in this study may be a consequence of differences in the levels or anatomical depth of transgene expression, rather than a reflection of their contribution to search behavior. Regardless of this experimental limitation, the fact that so many sensory modalities can trigger local searches suggests a convergence of these pathways onto the set of brain structures underlying navigation. This is consistent with anatomical studies of the central complex showing that it receives a variety of indirect sensory inputs [49], as well as direct innervation by a large subset of *NPF-GAL4* neurons [35, 54].

In summary, we found that hungry flies initiate a sustained local search when they encounter a fictive-food stimulus. This search behavior appears to be a generalized foraging response, as it can be triggered by multiple types of food-associated neurons including water-sensors, sugar-sensors, vinegar-odor-sensing neurons, as well as hunger-signaling neurons of the central nervous system. Like local searches triggered by real food, optogenetically-induced local searches are modulated by internal nutrient state and show key features of idiothetic path integration. Our results suggest that flies are able to keep track of their spatial position relative to a fictive food stimulus, even within a constrained maze. We demonstrate that long-lasting local search bouts can be initiated repeatedly by the brief activation of a small set of neurons, offering a promising entry-point to tracing the neural pathways underlying path integration in insects.

## ACKNOWLEDGEMENTS

We wish to thank Francesca V. Ponce for many helpful discussions about local search behavior, Rubi Salgado for help with fly rearing, Annie Rak for help with conducting the PER experiments, Ysabel M. Giraldo and Katherine J. Leitch for advice on statistical analysis, Floris van Breugel and Theodore H. Lindsay for assistance with implementing tracking software and closed-loop optogenetic stimulation, and William B. Dickson for assistance with image-processing for PER experiments. Kristin Scott and Hubert Amrein kindly provided us with flies. Research reported in this publication was supported by the National Institute of Neurological Disorders and Stroke of the National Institutes of Health under Award U19NS104655.

## METHODS

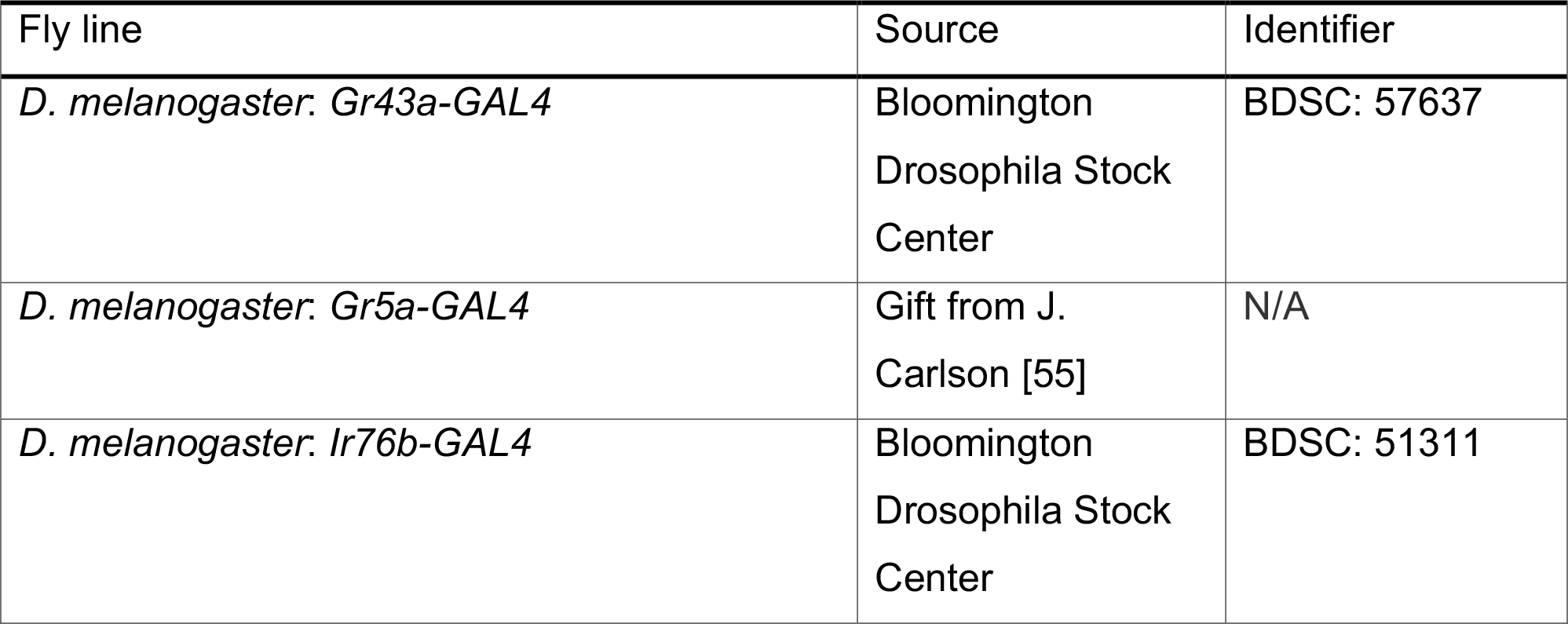

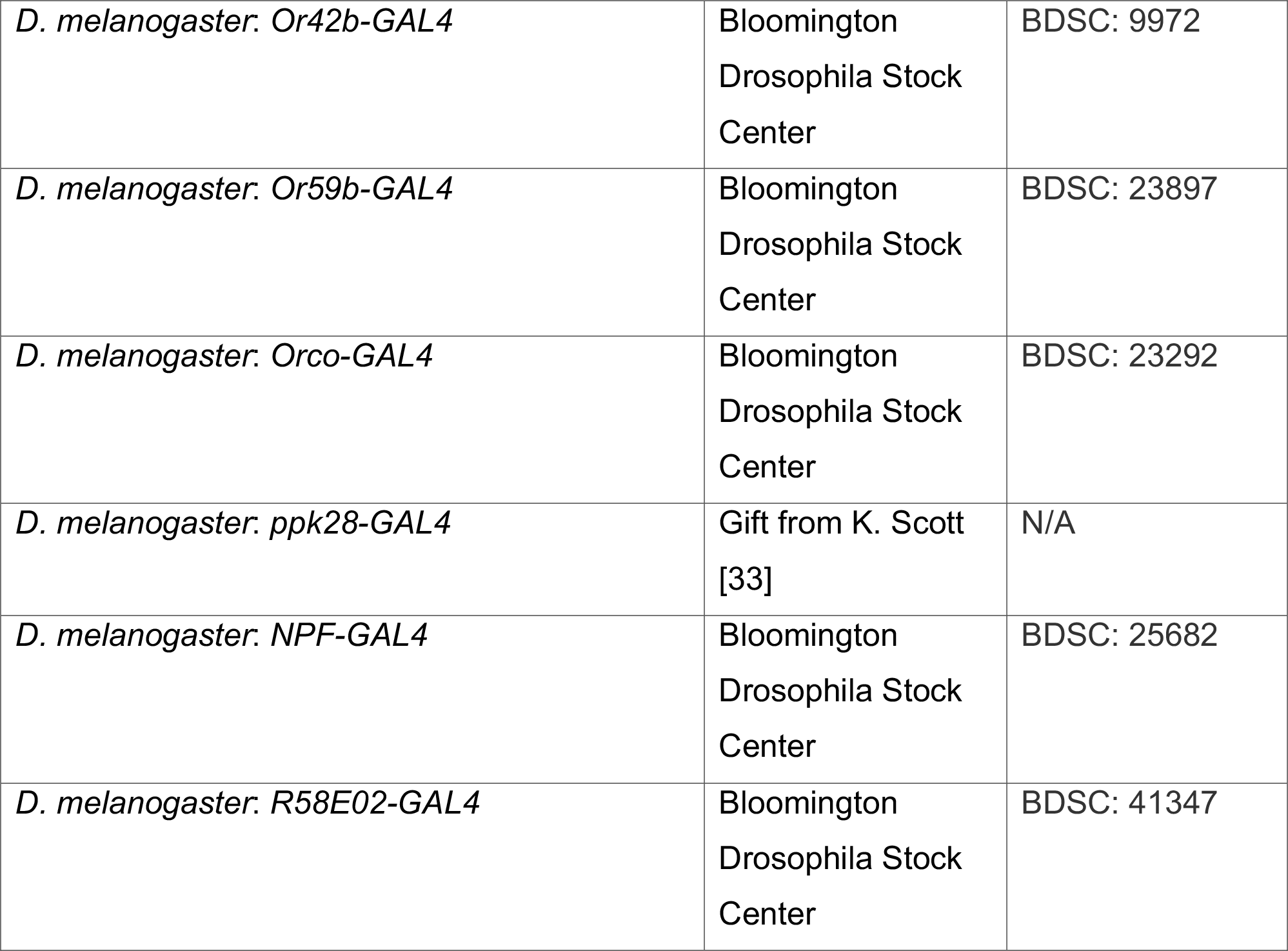

### Experimental animals

We conducted all experiments using 5-7-day old female *Drosophila melanogaster* reared in darkness at 25°C. Experimental flies were reared on standard cornmeal fly food containing 0.2 mM all transretinal (ATR) (Sigma) and transferred 0-2 days after eclosion onto standard cornmeal fly food with 0.4 mM ATR. Standard food was supplemented with additional yeast. Experimental flies were obtained by crossing *20XUAS-CsChrimson-mVenus* (inserted into attP40) virgin females to males of each GAL4 driver line. Parental control flies were obtained by crossing *20XUAS-CsChrimson-mVenus* (inserted into attP40) virgin females to wild-type males, and by crossing males of each GAL4 driver line to wild-type virgin females. The wild-type strain (Canton-S) originated form the lab of Martin Heisenberg.

Unless otherwise noted, we starved flies prior to experiments by housing them for 33-42 hours in a vial supplied with a tissue (KimTech) containing 1 ml of distilled water with 800 μM ATR. Protein-starved flies were housed for 5 days in a vial supplied with a tissue containing 1 mL of 10% sucrose with 800 μM ATR, and then transferred for 33-42 hours to a vial supplied with a tissue containing 1 ml of distilled water with 800 μM ATR. Dry-starved flies were housed for 16-26 hours in a vial containing a desiccant (Drierite) beneath a cotton ball. To facilitate sorting and transferring of experimental flies, we briefly anesthetized them at 4°C on a cold plate.

### Behavioral experiments

Experiments were conducted in a circular chamber constructed from layers of acrylic (118 mm diameter, 2.75 mm high). We constructed the flyadelphia chamber from layers of acrylic with insertable acrylic blocks to form the grid, such that flies were confined to the passages between blocks (1.85 mm wide, 2.75 mm high) but had sufficient space to walk forwards or backwards or turn around at any point in the arena. An upward-directed, custom-made array of 850 nm LEDs, covered by a translucent acrylic panel, was situated 11 cm beneath the arena to provide backlighting for a top-mounted camera (Blackfly, FLIR) recording at 30 frames per second. For optogenetic stimulation, we positioned upward-directed 628 nm LEDs (CP41B-RHS, Cree, Inc.) at the center of each activation zone, 8.5 mm beneath the arena floor. We covered the chamber lid with a long pass acrylic filter (color 3143, 3 mm thick). The chamber floor was transparent to allow for optogenetic stimulation, and a filter (#3000 Tough Rolux, Rosco Cinegel) was situated beneath the chamber to diffuse the red light used for stimulation. The camera, fly chamber, optogenetic stimulation lighting panel, and background lighting panel were held within a rigid aluminum frame (80/20), and the entire structure was covered with dense cloth to block any external light.

For each experiment, we aspirated a single fly into the behavioral chamber. Experiments began with an initial 10-minute baseline period, followed by 30 minutes during which the activation zone was operational. Experiments were terminated at the conclusion of this timeframe. Experiments in the flyadelphia arena consisted either of 60-minute trials during which the activation zone was operational, or equivalent baseline control trials with no activation zone. Behavioral chambers were cleaned with 100% ethanol at the conclusion of each trial and allowed to dry before reuse. We tracked the 2-D position of the fly in real-time using a python-based machine vision platform built on the Robot Operating System (http://florisvb.github.io/multi_tracker) [56]. The tracking system was customized to implement closed-loop control of optogenetic stimulation via an LED controller (1031, Phidgets Inc). The LED beneath each activation zone was turned on for 1 second whenever the centroid of the fly entered its virtual perimeter (diameter 5.67 mm), except during the baseline period. Each 1-second pulse was followed by a 9-second refractory period during which the LED was kept off, regardless of the fly’s position.

For PER experiments, we briefly anesthetized flies at 4°C on a cold plate and glued them to a tungsten wire at their dorsal surface anterior to the wings using UV-cured glue (Bondic, Inc.). We allowed tethered flies at least 30 minutes to recover, and then positioned flies with their head 1.3 mm from a fiber optic 617 nm light source for optogenetic stimulation. Flies were subjected to five 1-second light pulses, each separated by 20 seconds. Experiments were recorded via an overhead camera (Blackfly, FLIR, 30 frames per second). To determine the percentage of stimuli eliciting PER for each fly, we manually scored whether each light pulse elicited a proboscis extension during the 5 seconds following light pulse onset.

### Behavioral analysis

The dataset for each trial consisted of an array of X and Y coordinates representing the 2-D positions of the fly, as well as an array of LED states (on or off) for the activation zone(s). To process data, we discarded occasional frames where the fly was either not tracked, where a second object was tracked in addition to the fly (e.g. fly poop), or where the tracked jumped more than 1.5 mm within two consecutive frames (e.g. due to sporadic tracking of another object). Because the sampling rate of tracking data was not precisely 30 Hz, we down-sampled and linearly interpolated all data to 20 Hz to generate a regularly sampled time series.

We classified the fly as being within one of three possible locations in the arena: at the arena edge, in the open-field portion of the arena, or in the activation zone. The fly was considered to be at the arena edge if it was within 5.27 mm from the wall of the arena. For experiments in the flyadelphia arena, the arena edge was defined as all points more than 32.6 mm from the arena center. Otherwise, if the fly was not within the virtual perimeter of an activation zone, it was considered to be in the open-field portion of the arena. We used a Schmitt-trigger algorithm [57] to classify whether the fly was stopped or walking; a fly was considered to be stopped if its instantaneous speed fell below 1 mm s^−1^, or walking if its instantaneous speed rose above 3 mm s^−1^, and its classification state persisted when its speed fell between these thresholds.

We classified search bouts (Supplemental Figure 1A) as any trajectories beginning with a light pulse and ending when the fly reached the arena edge or at the conclusion of the trial. Equivalent bouts during baseline condition served as controls, and were defined as any trajectories beginning in the activation zone and ending when the fly reached the arena edge or at the conclusion of the trial. We divided search bouts into individual excursions from the activation zone, with each excursion counting as a revisit to the activation zone (Supplemental Figures 1B and 1C). To implement this, we defined two radial distances from the activation zone to serve as thresholds, again using the logic of a Schmitt trigger. Excursions began when a fly that had entered the activation zone crossed the excursion threshold ring in the outbound direction, and ended when the fly crossed the excursion threshold ring in the inbound direction (inner radius, 5.67 mm; outer radius, 7.09 mm). Throughout the paper, all analyses (unless otherwise noted) excluded search bouts truncated by the conclusion of the trial, as well as any timepoints when the fly was stopped or in the activation zone.

### Statistical analysis

We generated all figures using the python library FigureFirst (https://github.com/FlyRanch/figurefirst). Throughout the paper, 95% confidence intervals were calculated from distributions of median values generated by 1000 bootstrap iterations. Data replotted across figures may depict slightly differing 95% confidence intervals due to randomness inherent to the bootstrapping technique. Trials without a baseline control bout (i.e. the animal did not encounter the activation zone during the baseline period) were excluded for pair-wise Wilcoxon signed-rank tests.

---

**Figure S1.**
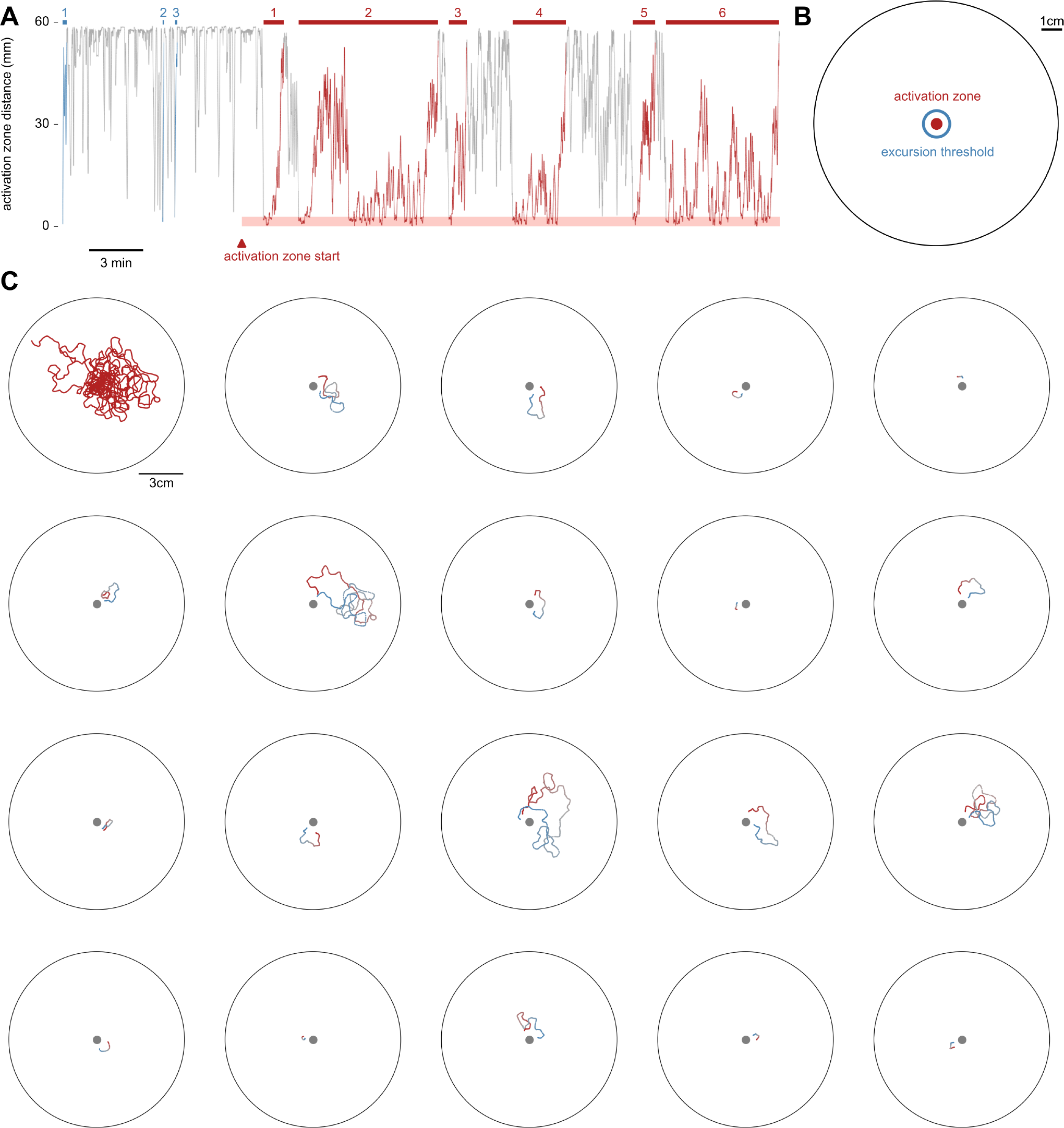
Local searches consist of excursions and revisits to the activation zone. (A) Example trial with a *Gr43a-GAL4>UAS-CsChrimson* animal, plotting distance between fly and activation zone center. Trials lasted 40 minutes, and the activation zone became operational after an initial 10-minute baseline control period. Red segments denote activation search bouts, defined as trajectories beginning with a light pulse and ending when the fly reached the arena edge or at the conclusion of the trial. Blue segments denote the equivalent baseline control bouts with no light pulse. (B) Schematic showing the 1.42 mm thick excursion threshold ring (blue) from the Schmitt-trigger analysis we used to divide search bouts into individual excursion events. An excursion begins when a fly that has received a light pulse crosses the excursion threshold ring in the outbound direction, and ends when the fly crosses the excursion threshold ring in the inbound direction. Each excursion is counted as a revisit to the activation zone. (C) Full trajectory (top left) of search bout number 6 from (A) and trajectories of all individual excursions therein. Excursion trajectories are color-coded from start (red) to end (blue).

**Figure S2.**
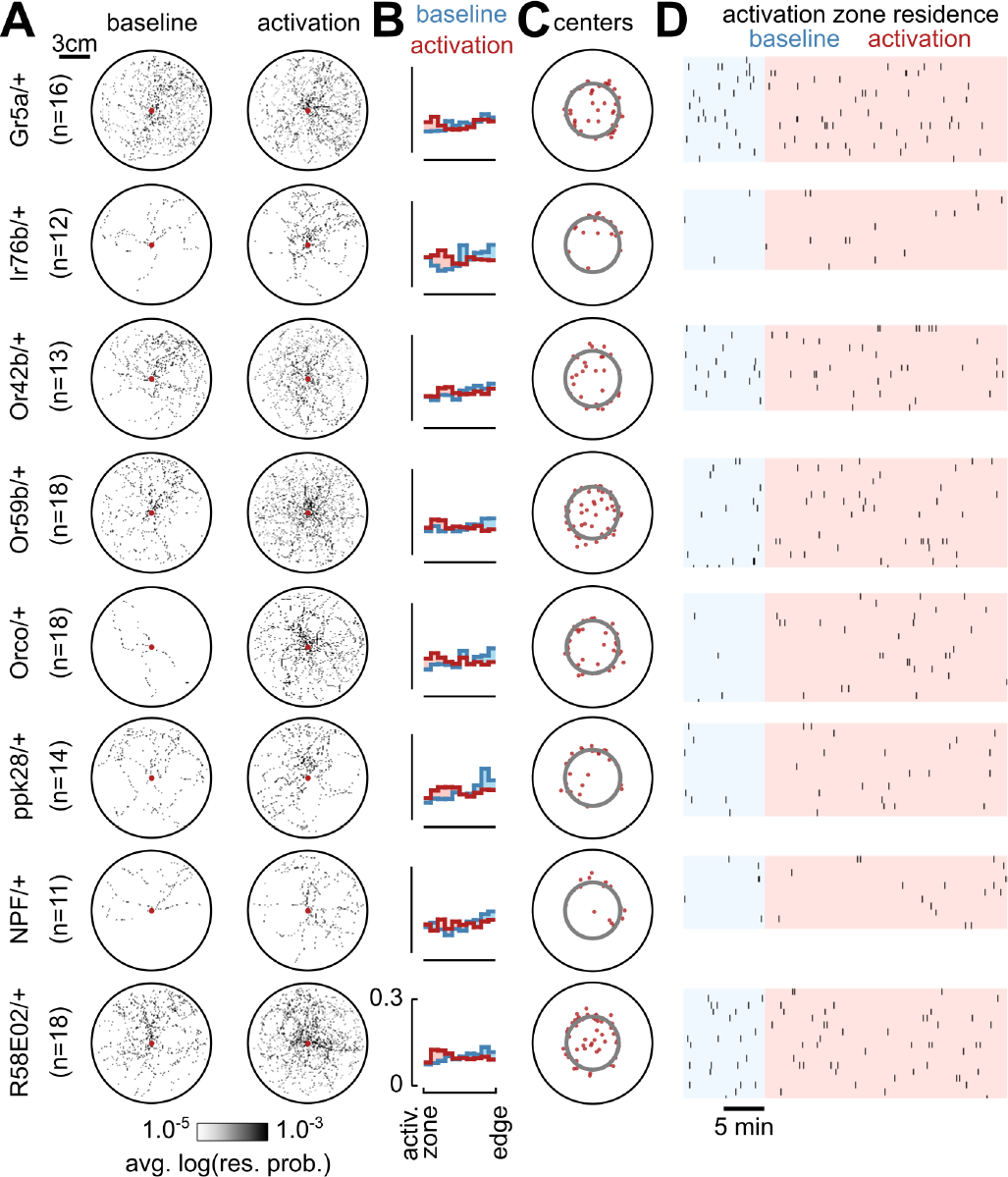
Additional parental controls for optogenetic screen. (A) Residence probabilities during activation search bouts and baseline control bouts, plotted as in Figure 1D. (B) Probability distributions of distance between fly and activation zone, plotted as in Figure 1E. (C) Centers of mass for all activation search bouts, plotted as in Figure 1F. (D) Raster plots of activation zone residence, plotted as in Figure 1G.

**Figure S3.**
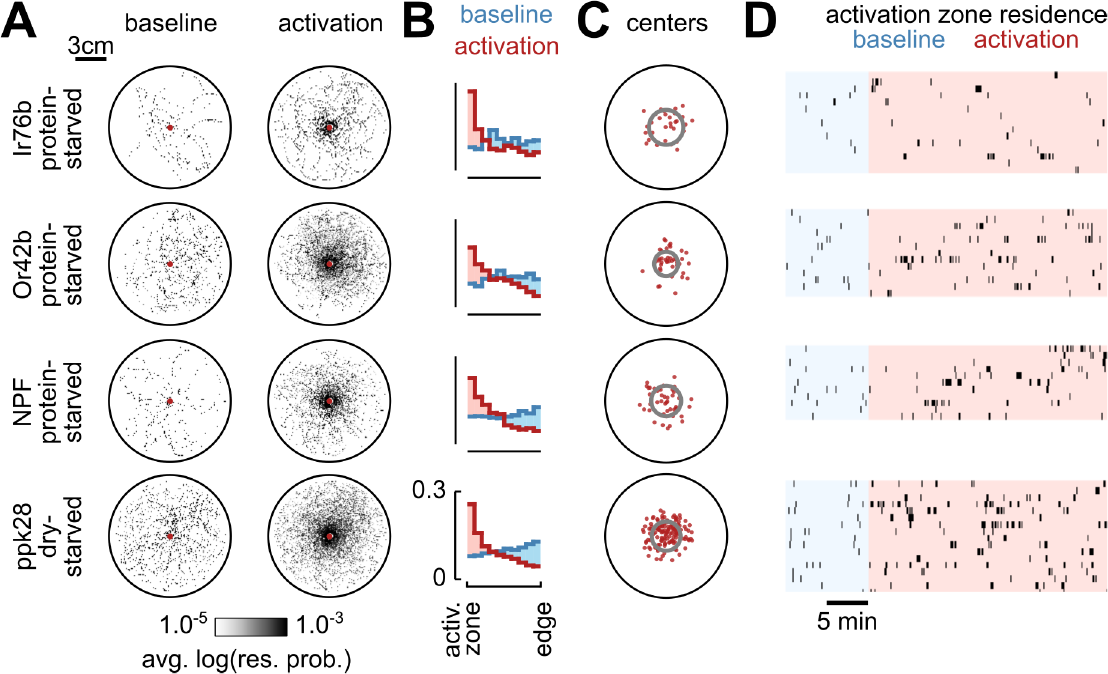
Additional analyses of local search in extreme-starved animals. (A) Residence probabilities during activation search bouts or baseline control bouts, plotted as in Figure 1D. Activation search bout residence probabilities reproduced from Figures 2G-J for convenience. (B) Probability distributions of fly distance to the activation zone, plotted as in Figure 1E. (C) Centers of mass for all activation search bouts, reproduced from Figures 2G-J for convenience. (D) Raster plots of activation zone residence, plotted as in Figure 1G.

**Figure S4.**
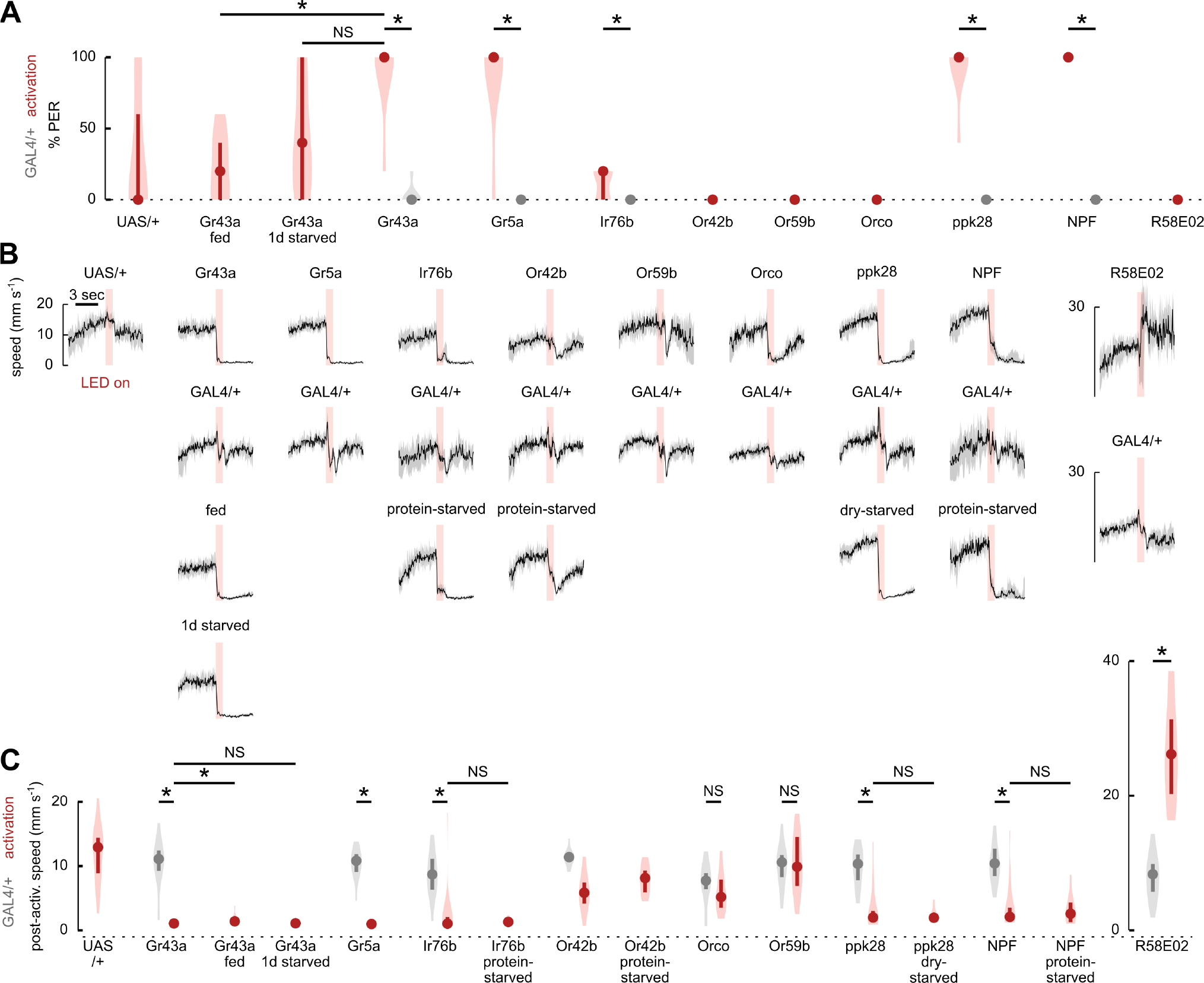
Proboscis extension and locomotor responses to activation. (A) Mean percentage of stimulus presentations eliciting a proboscis extension response (PER) for animals of the indicated genotype (red) or corresponding GAL4-line parental controls (grey) and indicated starvation state. Data depict distribution of individual fly means. Circles depict medians, error bars depict 95% confidence intervals, and violin plots indicate full data distribution. We note that light pulses produce PER in some *UAS-CsChrimson*/+ control animals, but we do not believe this hinders interpretation of results because we find no PER response in multiple other lines expressing this transgene. (* indicates p≤0.05, NS indicates p>0.05, n=7 flies per condition, Mann-Whitney U test with Bonferroni correction). (B) Median speed traces (solid black lines) during light pulse stimulation for animals of the indicated genotype and starvation state. Only the first light pulse for each search bout was used for analysis, and data were truncated if animals reached arena edge. Data depict distribution of individual fly median traces, with shaded grey region depicting 95% CIs. Data for *R58E02-GAL4>UAS-CsChrimson* animals are plotted on extended axes to accommodate values for higher speeds. (C) Median post-activation speed (during the 4 seconds following light pulse offset) for animals of the indicated genotype (red) or corresponding GAL4-line parental controls (grey) and indicated starvation state. Plotting conventions as in (A). Only the first light pulse for each search bout was used for analysis, and data were truncated if animal reached arena edge. Data for *R58E02-GAL4>UAS-CsChrimson* animals are plotted on extended axes to accommodate values for higher speeds.

**Movie 1. Activation of *Gr43a-GAL4* sugar-sensors triggers local search (raw footage).**

An example search bout triggered by activation of sugar-sensing *Gr43a-GAL4* neurons (corresponding to data shown in Figure1B). Approximately 19 seconds into the movie, the fly encounters the invisible activation zone (diameter 5.67 mm) at the center of the arena. The experiment was conducted in the dark. Footage was recorded using near-infrared (850 nm) lighting. The array of shadows visible in the movie were cast by the optogenetic activation LEDs and associated wiring situated beneath the chamber floor. Playback is at 5X speed.

**Movie 2. Activation of *Gr43a-GAL4* sugar-sensors triggers local search (animation).**

Animation of an example search bout triggered by activation of sugar-sensing *Gr43a-GAL4* neurons (corresponding to data shown in Figure1B and Movie 1). Fly position is marked by the green circle and the activation zone is shown as a red circle. Fly trajectory is shown before (blue) and after (red) the beginning of the search bout. LED pulses are indicated (bottom left). Playback is at 5X speed.

**Movie 3. Activation of *Gr5a-GAL4* sugar-sensors triggers local search.**

Animation of an example search bout triggered by activation of sugar-sensing *Gr5a-GAL4* neurons. Fly position is marked by the green circle and the activation zone is shown as a red circle. Fly trajectory is shown before (blue) and after (red) the beginning of the search bout. LED pulses are indicated (bottom left). Playback is at 5X speed.

**Movie 4. Activation of *Or42b-GAL4* olfactory neurons triggers local search in protein-starved animals.**

Animation of an example search bout triggered by activation of ACV-odor-sensing *Or42b-GAL4* neurons, in a protein-starved animal. Fly position is marked by the green circle and the activation zone is shown as a red circle. Fly trajectory is shown before (blue) and after (red) the beginning of the search bout. LED pulses are indicated (bottom left). Playback is at 5X speed.

**Movie 5. Activation of *NPF-GAL4* hunger-signaling neurons triggers local search in protein-starved animals.**

Animation of an example search bout triggered by activation of hunger-signaling *NPF-GAL4* neurons, in a protein-starved animal. Fly position is marked by the green circle and the activation zone is shown as a red circle. Fly trajectory is shown before (blue) and after (red) the beginning of the search bout. LED pulses are indicated (bottom left). Playback is at 5X speed.

**Movie 6. Activation of *ppk28-GAL4* water-sensing neurons triggers local search in dry-starved animals.**

Animation of an example search bout triggered by activation of water-sensing *ppk28-GAL4* neurons, in a dry-starved animal. Fly position is marked by the green circle and the activation zone is shown as a red circle. Fly trajectory is shown before (blue) and after (red) the beginning of the search bout. LED pulses are indicated (bottom left). Playback is at 5X speed.

**Movie 7. Optogenetically-induced local search in a confined maze.**

Animation of an example search bout in the flyadelphia arena, triggered by activation of sugar-sensing *Gr43a-GAL4* neurons. Fly position is marked by the green circle and the activation zone is shown as a red circle. Fly trajectory is shown before (blue) and after (red) the beginning of the search bout. LED pulses are indicated (bottom left). Playback is at 10X speed.

